# Activation of transient receptor potential vanilloid 4 is involved in pressure overload-induced cardiac hypertrophy

**DOI:** 10.1101/2021.10.25.465686

**Authors:** Yan Zou, Miaomiao Zhang, Qiongfeng Wu, Ning Zhao, Minwei Chen, Cui yang, Yimei Du, Bing Han

**Affiliations:** Department of Cardiology, Xuzhou Central Hospital, Xuzhou, Jiangsu 221009, China; Department of Cardiology, Union Hospital, Tongji Medical College, Huazhong University of Science and Technology, Wuhan, Hubei 430022, China; Department of Cardiology, Xiamen Key Laboratory of Cardiac Electrophysiology, Xiamen Institute of Cardiovascular Diseases, The First Affiliated Hospital of Xiamen University, School of Medicine, Xiamen University, Xiamen 361003, China; Xuzhou Institute of Cardiovascular Disease, Xuzhou Central Hospital, Xuzhou, Jiangsu 221009, China.

**Keywords:** TRPV4, Ca^2+^/calmodulin-dependent protein kinase II, NFκB, NOD-like receptor pyrin domain-containing protein 3, cardiac hypertrophy, mechanosensitive channels, medicine, mouse.

## Abstract

Previous studies, including our own, have demonstrated that transient receptor potential vanilloid 4 (TRPV4) is expressed in hearts and implicated in cardiac remodeling and cardiac dysfunction. However, the effects of TRPV4 on pressure overload-induced cardiac hypertrophy remain unclear. In this study, we found that TRPV4 expression was significantly increased in mouse hypertrophic hearts, human failing hearts, and neurohormone-induced hypertrophic cardiomyocytes. Deletion of TRPV4 attenuated transverse aortic constriction (TAC)-induced cardiac hypertrophy, cardiac dysfunction, fibrosis, inflammation, and the activation of NFκB - NOD-like receptor pyrin domain-containing protein 3 (NLRP3) in mice. In vitro, TRPV4 inhibition decreased the neurohormone-induced cardiomyocyte hypertrophy and the increase of intracellular Ca^2+^ concentration. TRPV4 agonist triggered Ca^2+^ influx and evoked the phosphorylation of Ca^2+^/calmodulin-dependent protein kinase II (CaMKII) but these effects were abolished by removing extracellular Ca^2+^ or TRPV4 inhibition. More importantly, TAC or neurohormone stimulation-induced CaMKII phosphorylation was significantly blocked by TRPV4 inhibition. Finally, we showed that CaMKII inhibition significantly inhibited the phosphorylation of NFκB induced by TRPV4 activation. Our results suggest that TRPV4 activation contributed to pressure overload-induced cardiac hypertrophy. This effect was associated with upregulated Ca^2+^/ CaMKII mediated the activation of NFκB-NLRP3. Thus, TRPV4 may represent a potential therapeutic drug target for cardiac hypertrophy.

## Introduction

In response to pathological stimuli such as hypertension, valvular heart disease, or neurohumoral overactivation, the heart undergoes hypertrophy. Initially, the hypertrophy response is adaptive; however, sustained cardiac hypertrophy results in increased heart mass, cardiac fibrosis, and eventually heart failure(Bui, et al., 2011; Nakamura and Sadoshima, 2018). Although significant advances in the treatment of pathological hypertrophy, heart failure still is a leading cause of death worldwide(Neubauer, 2007). Thus, to deeply uncover the molecular mechanism of pathological cardiac hypertrophy continues to be important for developing novel therapeutic strategies for the prevention of cardiac remodeling and dysfunction(Kalman, et al., 2019).

Increased mechanical stress plays a key role in cardiac hypertrophy. The transient receptor potential vanilloid (TRPV) channels are ubiquitous ion channels that function as essential mechanical sensors(Clapham, 2003). Interestingly, those channels are upregulated in the hearts of mice after transverse aortic constriction (TAC), as shown for TRPV1, TRPV2, and TRPV3(Chen, et al., 2016; Lorin, et al., 2015; Zhang, et al., 2018). Furthermore, the genetic deletion of functional TRPV2 ameliorates significantly TAC-induced cardiac hypertrophy and dysfunction(Koch, et al., 2017). These findings suggest the critical role of TRPV in the development of cardiac remodeling in response to pressure overload.

TRPV4, a member of the TRPV subfamily, is wildly expressed in the cardiovascular system(Hof, et al., 2019; White, et al., 2016). Its functional expression is increased under certain pathological conditions, such as pressure overload(Morine, et al., 2016), aging(Jones, et al., 2019), ischemia-reperfusion(Dong, et al., 2017; Wu, et al., 2017), and pericarditis(Liao, et al., 2020). Inhibition of TRPV4 attenuates intracellular calcium concentration ([Ca^2+^]i)(Wu, et al., 2017), cardiac fibrosis(Adapala, et al., 2020), and cardiac inflammation(Liao, et al., 2020), which improves cardiac function(Wu, et al., 2019). In addition, a potent and selective TRPV4 inhibitor recently revealed a positive efficacy trend in a phase 2a trial in patients with heart failure(Goyal, et al., 2019; Stewart, et al., 2020). To date, TRPV4 has not been reported in association with pressure overload-induced cardiac hypertrophy. Therefore, in the present study, we aimed to investigate the role and the underlying mechanism of TRPV4 in pathological cardiac hypertrophy subjected to pressure overload.

## Results

### TRPV4 expression is increased in pathological cardiac hypertrophy

To evaluate the potential role of TRPV4 in cardiac hypertrophy, we firstly measured TRPV4 protein and mRNA expression levels in left ventricle (LV) tissue from wild-type (WT) TAC vs. sham mice 4 weeks after surgery. As shown in Figures 1A-B, TAC induced a two-fold increase in TRPV4 protein expression. This finding was confirmed with a two-fold increase in mRNA expression in wild-type TAC hearts (Figure 1C). We also assessed the TRPV4 expression level in LV tissue from human hearts and found that TRPV4 protein was significantly upregulated in failing hearts compared with non-failing (Figures 1D-E). Our results indicate that TRPV4 may be implicated in the development of pathological cardiac hypertrophy.

**Figure 1.**
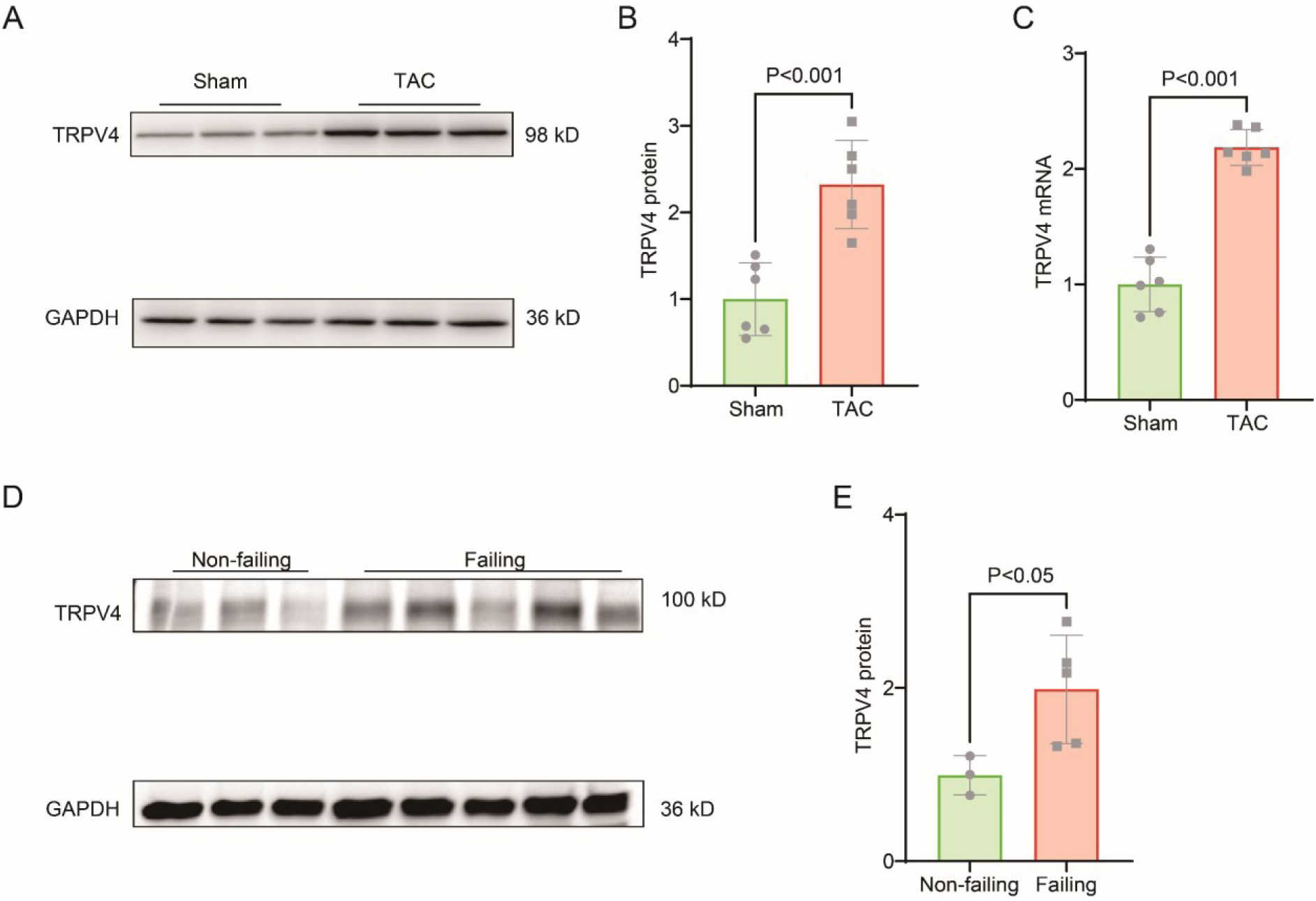
TRPV4 expression is upregulated in pathological cardiac hypertrophy. Representative immunoblot image (**A**) and statistics (**B**) of TRPV4 protein level in the LV from sham or TAC mice (n = 6 per group). **C.** Statistical data of TRPV4 mRNA level in the LV from sham or TAC mice (n = 6 per group). Representative immunoblot image (**D**) and statistical data (**E**) of TRPV4 protein level in human non-failing hearts (n = 3) and failing hearts (n = 5). All results represent mean ± SD, an unpaired two-tailed student’s t-test.

### TRPV4 deficiency attenuates cardiac hypertrophy induced by pressure overload in vivo

To further investigate the role of TRPV4 in cardiac hypertrophy induced by pressure overload, we performed TAC or sham surgery in WT and TRPV4 knock-out (TRPV4 -/-) mice (Figure 2A). We used the ratios of heart weight/body weight (HW/BW) and heart weight /tibial length (HW/TL) to assess changes in LV mass (Figure 2B-C). As expected, both values in sham-operated WT and TRPV4 mice were similar (HW/BW ratio: 4.49 ± 0.09 vs 4.63 ± 0.13, HW/TL ratio: 6.34 ± 0.19 vs 6.83 ± 0.18). TAC induced a 59 and 52% increase (all *P*<0.001) in HW/BW ratio and HW/TL ratio, respectively, in WT mice. However, this hypertrophic response to TAC was attenuated in TRPV4-/-mice, as evident by only an 18 and 12% increase in HW/BW ratio (*P*<0.01) and HW/TL ratio (*P* <0.05), respectively. Next, we measured the cross-sectional area of myocytes in all groups. As shown in Figure 2D, cell surface area increased in both WT and TRPV4-/- mice after TAC, however, the increase was significantly attenuated in TRPV4-/- myocytes compared with WT (281.25±39.69 μm^2^ vs 547.17±109.26 μm^2^, *P*<0.001). In order to confirm our findings at the molecular level, we then detected cardiac hypertrophic marker genes expression. Both ANP and BNP mRNA expression were significantly higher in WT hearts compared with TRPV4-/- hearts after TAC. There was no significant difference between WT and TRPV4-/- in the sham group (Figure 2E-F). These results suggest that TRPV4 activation plays a critical role in pressure overload-induced cardiac hypertrophy.

**Figure 2.**
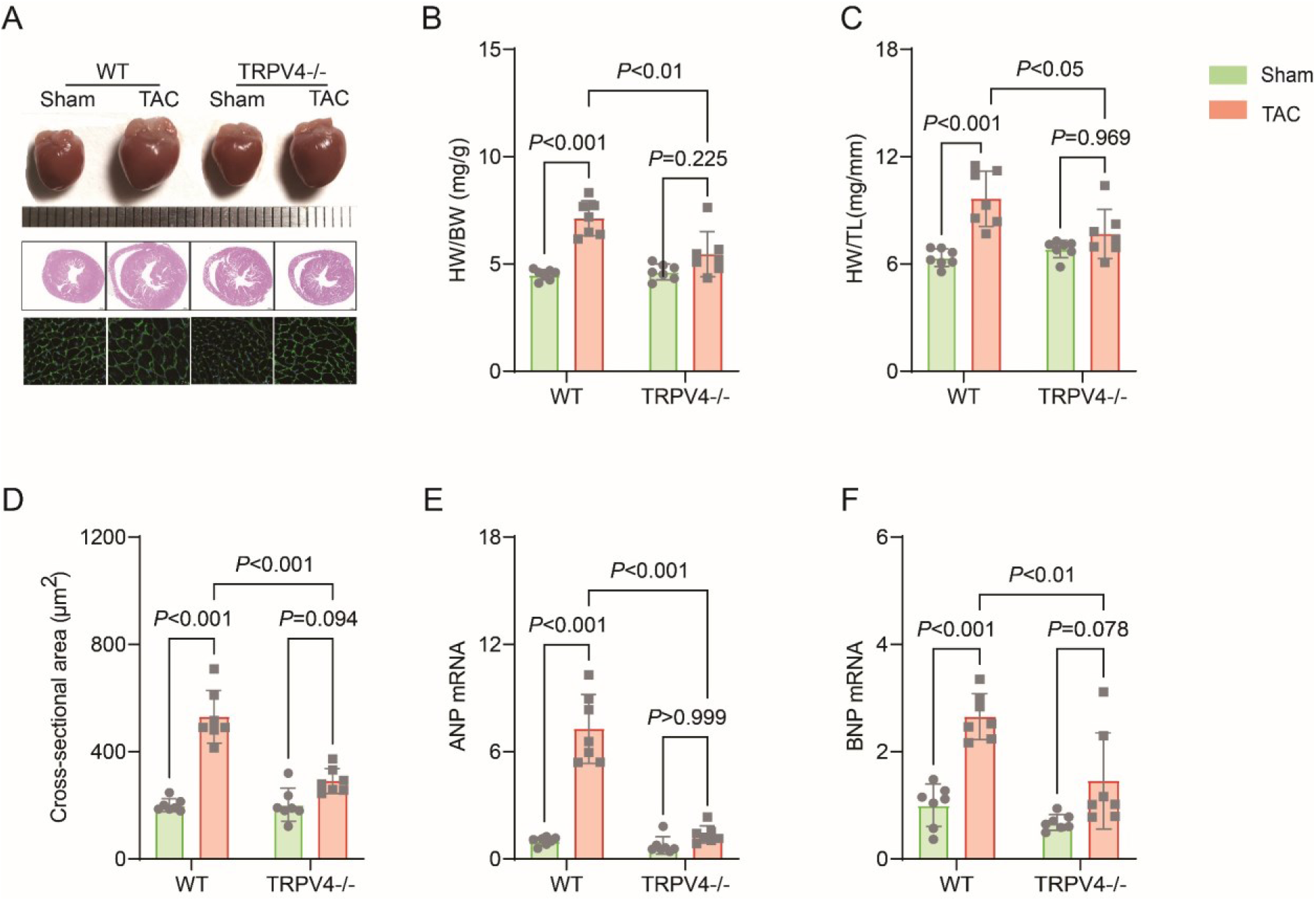
TRPV4 deficiency attenuates pressure overload-induced cardiac hypertrophy. Representative images of heart photo, H&E staining, and WGA staining of WT and TRPV4-/- mice at 4 weeks after sham or TAC operation (**A**). Statistical results for the ratios of HW/BW (**B**), HW/TL (**C**), and cross-section area (D) in mice 4 weeks after sham or TAC operation (n = 7 per group). Statistics of hypertrophy-related genes ANP (**E**) and BNP (**F**) mRNA levels in mouse hearts 4 weeks after sham or TAC operation (n = 7 per group). All results represent mean ± SD, a two-way ANOVA followed by the Bonferroni test.

### TRPV4 deficiency attenuates cardiac dysfunction and cardiac fibrosis induced by pressure overload

Echocardiography was performed to monitor the progression of cardiac structure and functional changes (Figure 3A). A reduction in ejection fraction (EF, 52.83±4.34% vs 73.44±2.47%, *P*<0.001, Figure 3B) and fractional shortening (FS, 26.87±2.64% vs 41.34±1.97%, *P*<0.001, Figure 3C) in WT mice were reversed in TRPV4-/- mice at 4 weeks after TAC. Consistently, LV internal dimension systole and LV mass were significantly increased in WT TAC mice, but these effects were not found in TRPV4-/- TAC mice (Figured 3D-E). Other parameters of LV remodeling including LV posterior end-diastolic wall thickness (LVPW), LV end-diastolic diameter (LVEDD), and LV end-diastolic volume (LVEDV) were also well preserved in TRPV4-/- mice compared with WT mice after TAC (Table 1).

**Figure 3.**
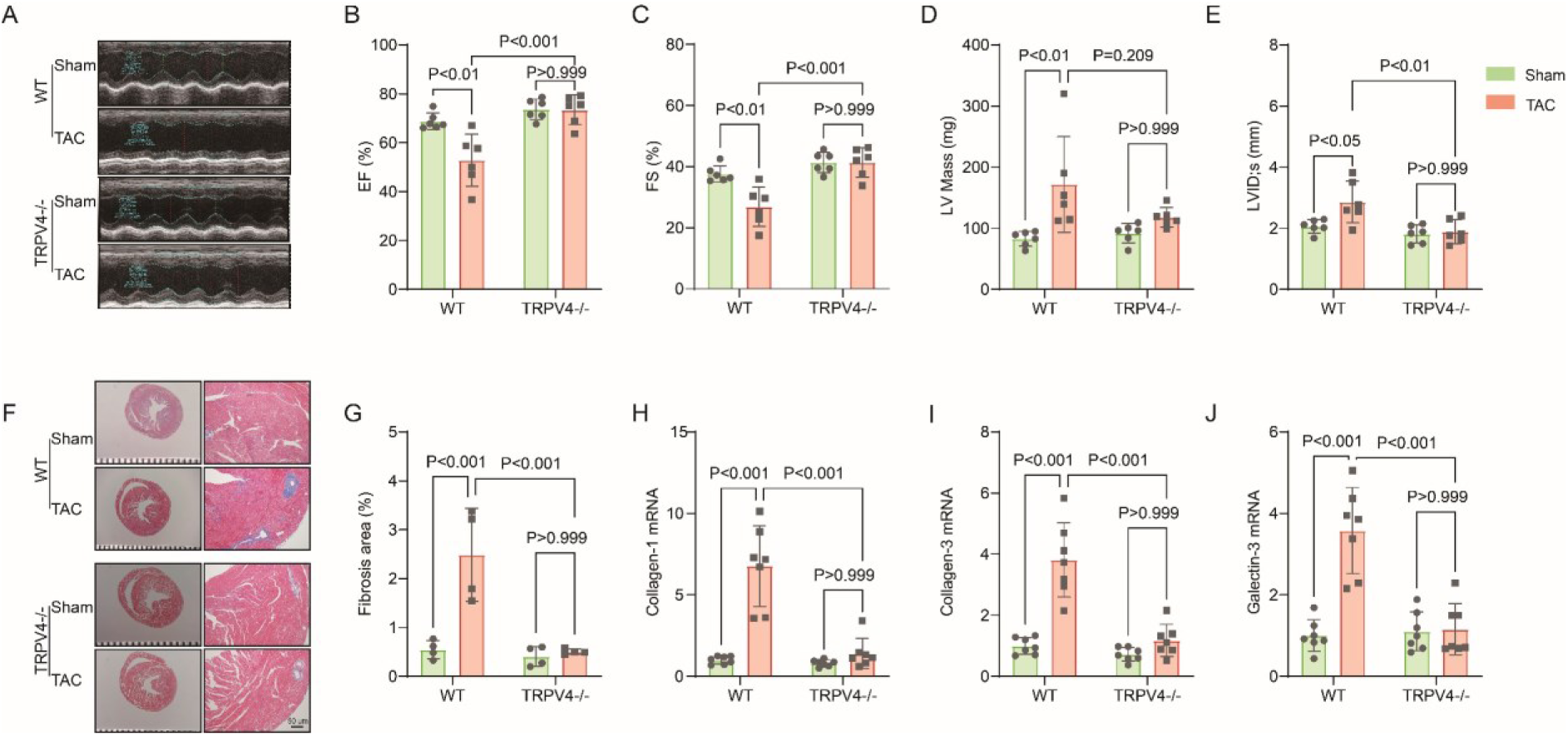
TRPV4 deficiency improves cardiac function and attenuates cardiac fibrosis induced by pressure overload. Representative images of M-mode echocardiography of WT and TRPV4-/- mice at 4 weeks after sham or TAC operation (**A**). Statistics of EF (**B**), FS (**C**), LV mass (**D**), and LVIDs (**E**) in mice at 4 weeks after sham or TAC operation (n = 6 per group). Representative images (**F**) and statistics (**G**) of Masson’s trichrome-stained hearts from mice at 4 weeks after sham or TAC operation. The statistics were from the panoramic scanning pictures (n = 4 each group). Statistics of fibrosis-related genes collagenase-1 (**H**), collagenase- 3 (**I**), and galectin-3 (**J**) mRNA levels in mouse hearts 4 weeks after sham or TAC operation (n = 7 per group). All results represent mean ± SD, a two-way ANOVA followed by the Bonferroni test.

**Table 1.**
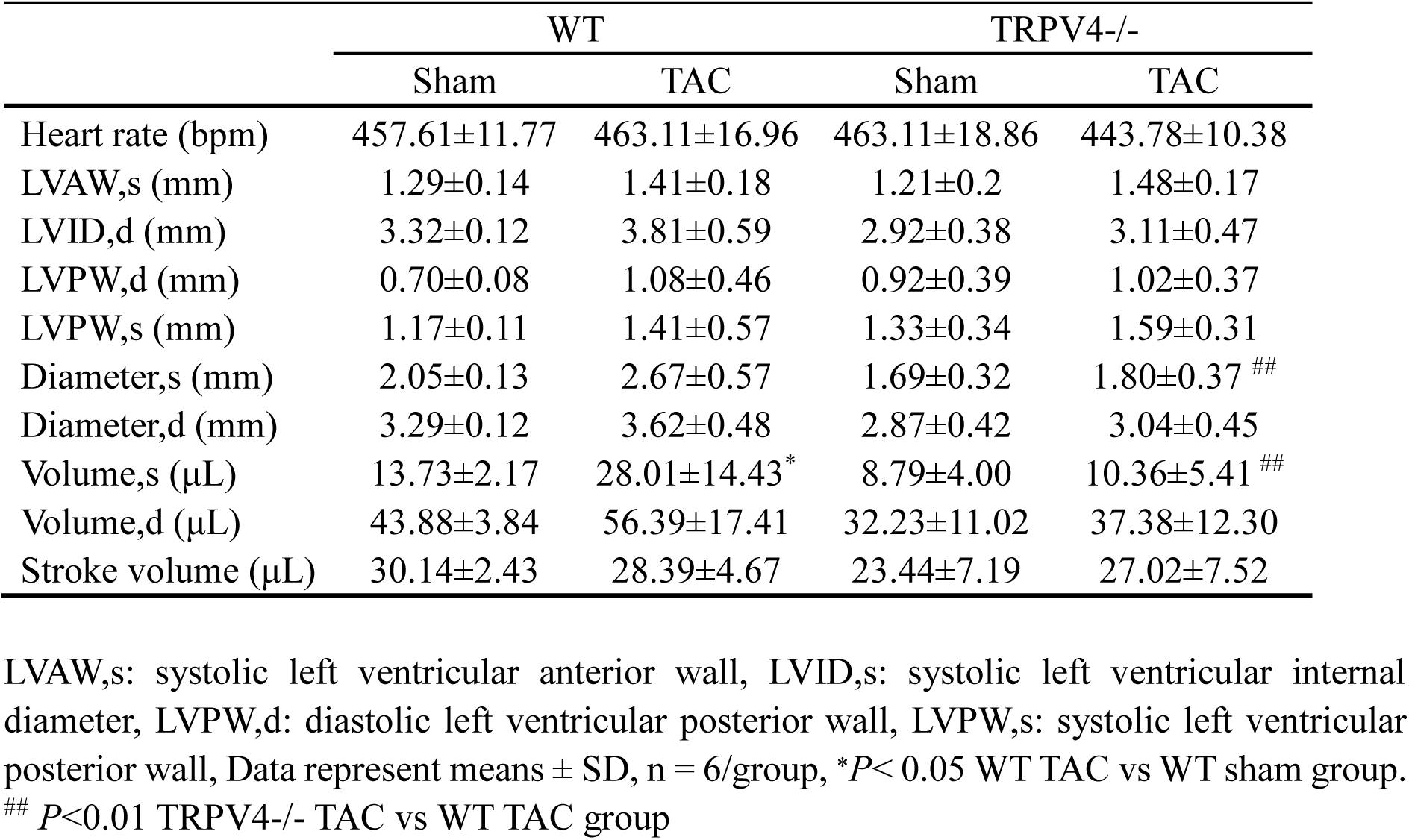
Echocardiographic measurements 4 weeks after TAC

|  | WT |  | TRPV4<sup>-/-</sup> |  |
| --- | --- | --- | --- | --- |
|  | Sham | TAC | Sham | TAC |
| Heart rate (bpm) | 457.61±11.77 | 463.11±16.96 | 463.11±18.86 | 443.78±10.38 |
| LVAW,s (mm) | 1.29±0.14 | 1.41±0.18 | 1.21±0.2 | 1.48±0.17 |
| LVID,d (mm) | 3.32±0.12 | 3.81±0.59 | 2.92±0.38 | 3.11±0.47 |
| LVPW,d (mm) | 0.70±0.08 | 1.08±0.46 | 0.92±0.39 | 1.02±0.37 |
| LVPW,s (mm) | 1.17±0.11 | 1.41±0.57 | 1.33±0.34 | 1.59±0.31 |
| Diameter,s (mm) | 2.05±0.13 | 2.67±0.57 | 1.69±0.32 | 1.80±0.37<sup>##</sup> |
| Diameter,d (mm) | 3.29±0.12 | 3.62±0.48 | 2.87±0.42 | 3.04±0.45 |
| Volume,s (μL) | 13.73±2.17 | 28.01±14.43<sup>*</sup> | 8.79±4.00 | 10.36±5.41<sup>##</sup> |
| Volume,d (μL) | 43.88±3.84 | 56.39±17.41 | 32.23±11.02 | 37.38±12.30 |
| Stroke volume (μL) | 30.14±2.43 | 28.39±4.67 | 23.44±7.19 | 27.02±7.52 |

LVAW,s: systolic left ventricular anterior wall, LVID,s: systolic left ventricular internal diameter, LVPW,d: diastolic left ventricular posterior wall, LVPW,s: systolic left ventricular posterior wall, Data represent means ± SD, n = 6/group, <sup>\*</sup>*P* < 0.05 WT TAC vs WT sham group. <sup>##</sup> *P* < 0.01 TRPV4<sup>-/-</sup> TAC vs WT TAC group

Cardiac interstitial and perivascular fibrosis were assessed in Masson’s Trichrome stained sections at 4 weeks after TAC surgery (Figure 3F). There was no significant difference in the extents of fibrosis in WT and TRPV4-/- mice in the sham groups. However, both interstitial and perivascular fibrosis increased in WT hearts after TAC, with more pronounced perivascular changes. This increase in interstitial and perivascular fibrosis was significantly blunted in TRPV4-/- hearts after TAC (2.48±0.95% vs 0.41±0.20%, *P*<0.001, Figure 3G). In addition, quantitative real-time PCR revealed a marked reduction in fibrosis markers (collagenase-1, collagenase-3, and galectin-3, Figure 3H-J).

### TRPV4 deficiency attenuates the inflammation induced by pressure overload

Chronic inflammation promotes cardiac fibrosis(Adamo, et al., 2020). Thus, we detected the protein and mRNA levels of pro-inflammatory cytokines. As shown in Figures 4A-D, TAC significantly upregulated the protein levels of IL-1β, IL-6, and TNF-α in WT mice, and TRPV4 deletion diminished this elevation. Consistent with these observations, the TAC-induced increases in mRNA expression of IL-1β, IL-6, TNF-α, MIP-2, and MCP-1 were significantly attenuated in TRPV4-/- mice (Figures 4E-I).

**Figure 4.**
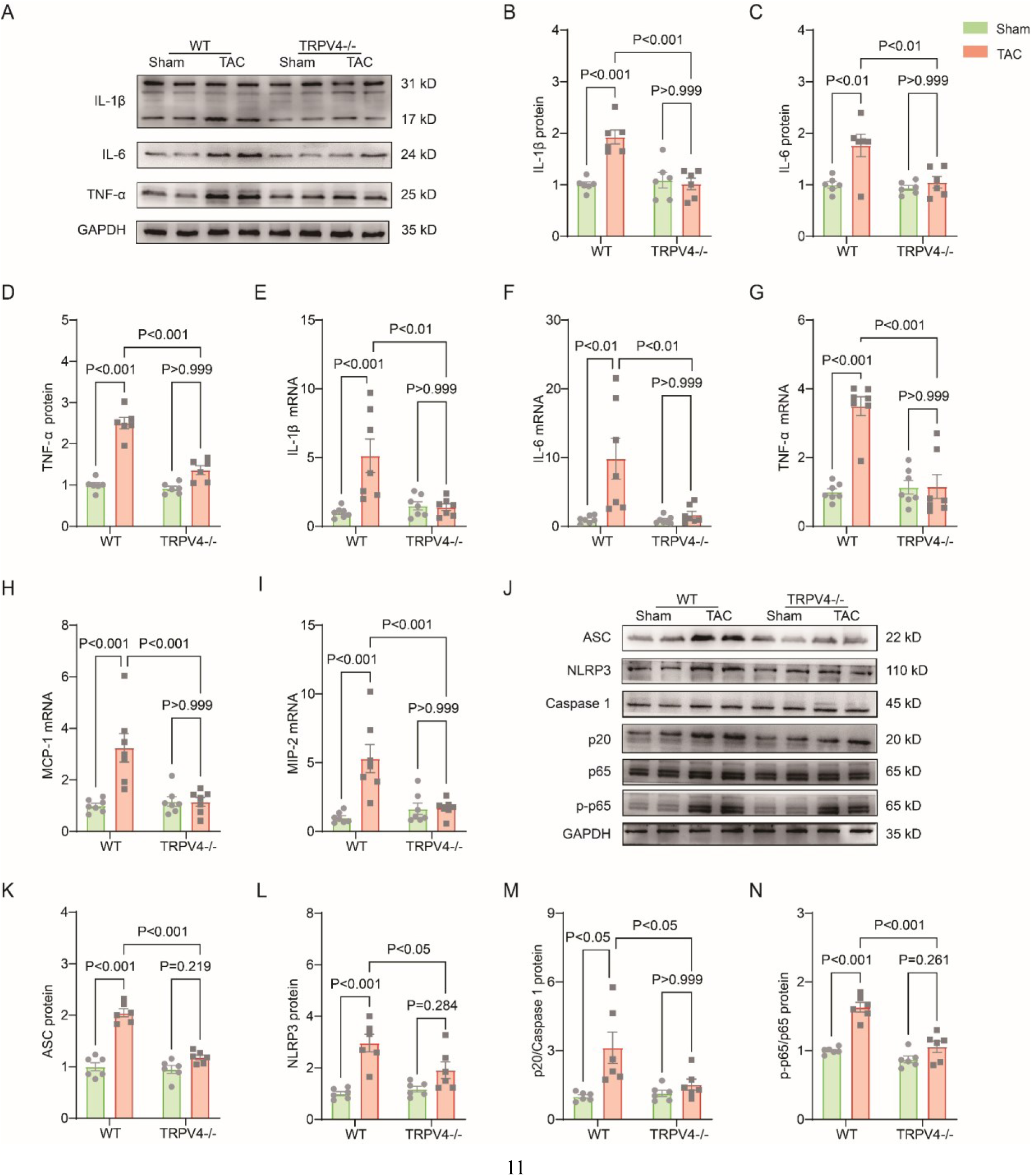
TRPV4 deficiency attenuates cardiac fibrosis induced by pressure overload. Representative immunoblot image (**A**) and statistics of IL-1β (**B**), IL-6 (**C**), and TNF-α (**D**) protein levels in WT and TRPV4-/- mice at 4 weeks after sham or TAC operation (n = 6 per group). Statistical data of IL-1β (**E**), IL-6 (**F**), TNF-α (**G**), MCP-1 (**H**), and MIP-2 (**I**) mRNA levels in mouse hearts 4 weeks after sham or TAC operation (n = 7 per group). Representative immunoblot image (**J**) and statistics of ASC (**K**), NLRP3 (**L**), Caspase 1-p20 (**M**), and p-NFκB p65 (**N**) protein levels in WT and TRPV4-/- mice at 4 weeks after sham or TAC operation (n = 6 per group). All results represent mean ± SD, a two-way ANOVA followed by the Bonferroni test.

The NOD-like receptor pyrin domain-containing protein 3 (NLRP3) inflammasome consists of ASC, NLRP3, and caspase-1(Martinon and Tschopp, 2004). Its activation contributes to the development of cardiac hypertrophy by cleaving pro-caspase-1 and promoting the release of proinflammatory cytokine IL-β(Suetomi, et al., 2019; Suetomi, et al., 2018). NFκB represents a family of inducible transcription factors, which regulates various genes involved in inflammatory responses. We then assessed the activation of NLRP3 inflammasome and the phosphorylation of NFκB (Figure 4J). As shown in Figures 4K-L, TAC significantly upregulated the protein levels of ASC, NLRP3, and cleaved caspase-1 (p20) in WT mice. We also found the expression of p-NFκB p65 (ser536) was greatly upregulated in WT mice after TAC surgery (Figure 4N). Interestingly, TRPV4 deletion efficiently decreased the ASC, NLRP3, cleaved caspase-1, and p-NFκB p65 protein levels.

### The TRPV4 antagonist improves neonatal rat ventricular myocytes (NRVMs) hypertrophy in vitro

Next, we sought to determine whether TRPV4 activation contributes to cardiomyocyte hypertrophy in vitro. NRVMs were isolated from neonatal Sprague-Dawley (SD) rats and treated with angiotensin II (Ang II) or phenylephrine (PE) for 48 hours. We found that AngII-stimulated cardiac hypertrophy, as indicated by increases in cell surface area (Figures 5 A-B) and expression of ANP and BNP (Figures 5 C-D), were largely inhibited by the TRPV4 specific antagonist GSK2193874 (GSK3874). Similarly, PE-induced CM hypertrophy was also attenuated by GSK3874 (Figures 5E-H). Taken together, our results confirmed that TRPV4 activation contributes to cardiac hypertrophy in vitro.

**Figure 5.**
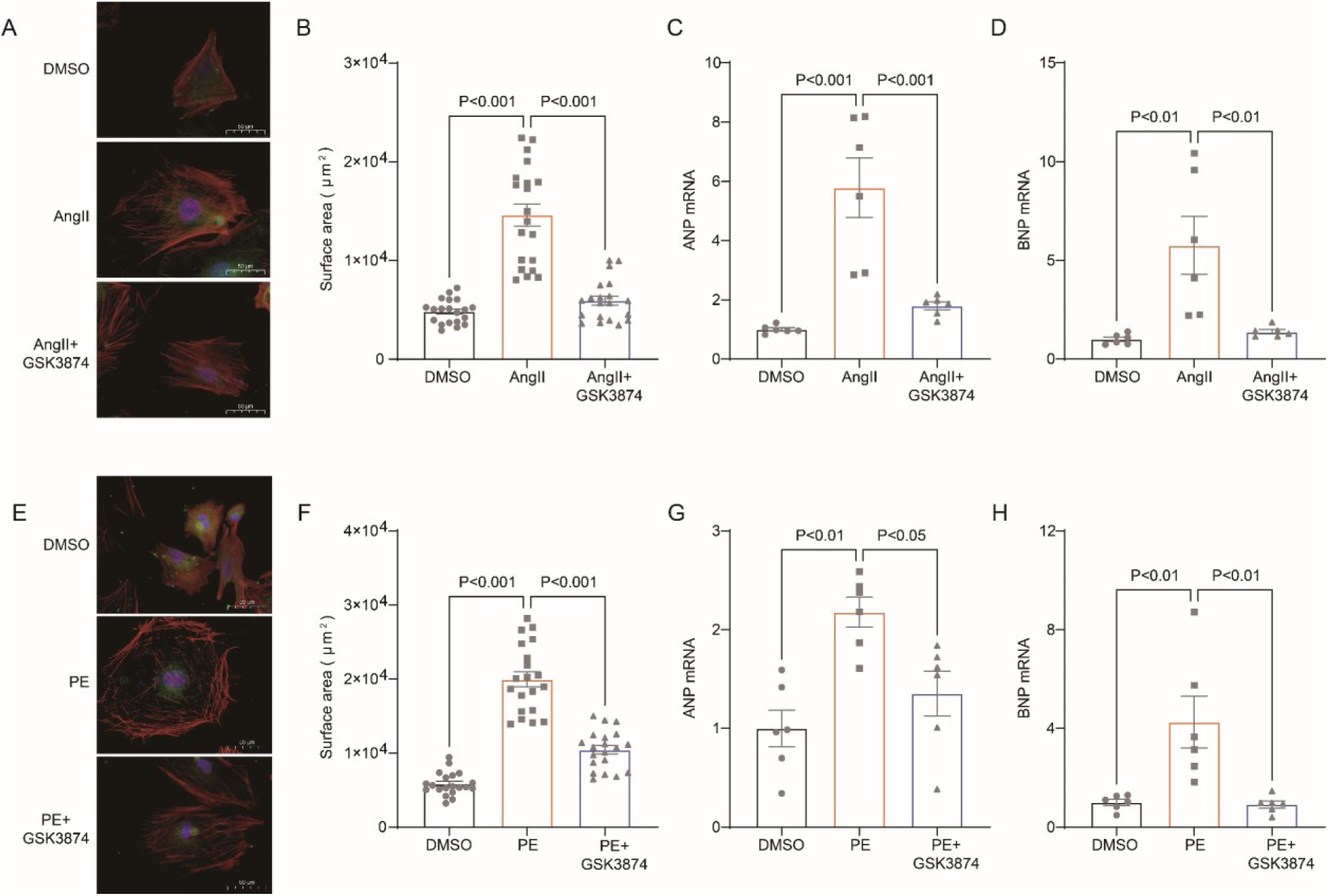
TRPV4 blockade attenuates AngII/PE-induced hypertrophy in NRVMs *in vitro*. Representative images (**A**) and statistics of the cell-surface areas (**B**) in NRVMs treated with DMSO, AngII, and AngII plus GSK3874 (n = 20 cells from 3 animals. Statistics of ANP (**C**) and BNP (**D**) mRNA levels in NRVMs treated with DMSO, AngII, and AngII plus GSK3874 (n = 6 per group). Representative images (**E**) and statistics of the cell-surface areas (**F**) in NRVMs treated with DMSO, PE, and PE plus GSK3874 (n = 20 cells from 3 animals). Statistics of ANP (**G**) and BNP (**H**) mRNA levels in NRVMs treated with DMSO, PE, and PE plus GSK3874 (n = 6 per group). All results represent mean ± SD, a one-way ANOVA followed by the Bonferroni test.

### TRPV4 antagonist alleviates on AngII/PE induced Ca^2+^ overload in NRVMs

It is well known that the [Ca^2+^]i increases in response to sustained hypertrophy. We have previously shown that TRPV4 functionally expresses in cardiomyocytes and mediates Ca^2+^ influx upon activation(Wu, et al., 2017). Here, we found that TRPV4 protein and mRNA expression was significantly increased in NRVMs after being treated with AngII (Figure 6A-C). To correlate TRPV4 expression to functional channel, changes in [Ca^2+^]i in response to the specific TRPV4 agonist GSK1016790A (GSK790A, 500 nM), were measured in NRVMs after AngII stimulation. As shown in Figure 6D-E, GSK790A induced robust Ca^2+^ influx, which was further enhanced after stimulation with AngII. However, pre-incubation of GSK3874 could inhibit this enhanced response. Please note, treatment with AngII or AngII + GSK3874 had no effect on Ca^2+^ influx induced by A23187 (Figure 6F). Similar results were also obtained from NRVMs after PE stimulation (Figures 6G-I). Our results indicate that TRPV4 activation may be implicated in [Ca^2+^]i rise induced by sustained hypertrophy.

**Figure 6.**
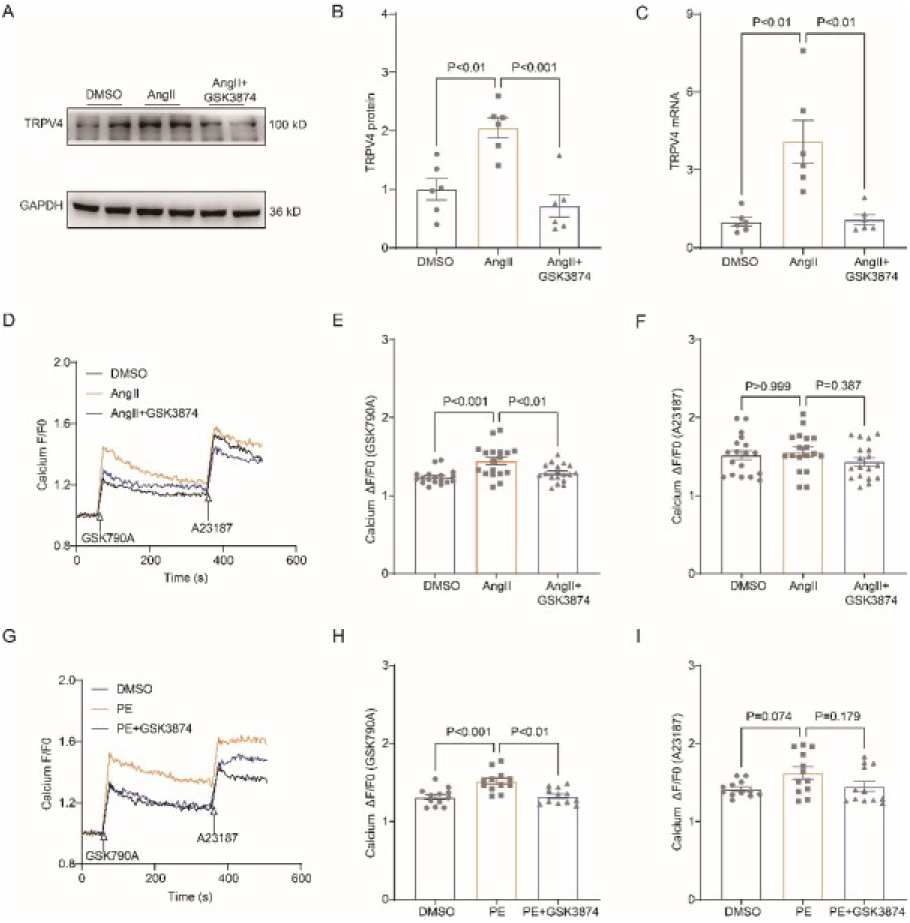
TRPV4 blockade attenuates AngII/PE-induced Ca^2+^ overload in NRVMs. Representative immunoblot image (**A**) and statistics (**B**) of TRPV4 protein level in NRVMs treated with DMSO, AngII, and AngII plus GSK3874 (n = 6 per group). **C**. Statistical data of TRPV4 mRNA level in NRVMs treated with DMSO, AngII, and AngII plus GSK3874 (n = 6 per group). Representative recording of changes in intracellular Ca^2+^ induced by 500 nM GSK 790A and 1 μM A23187 in NRVMs treated with DMSO, AngII, and AngII plus GSK3874 (**D**). Quantification of [Ca^2+^]i response induced by GSK790A (**E**) and A23187(**F**) -induced in NRVMs treated with DMSO, AngII, and AngII plus GSK3874 (n = 18 per group). Representative recording of changes in intracellular Ca^2+^ induced by 500 nM GSK 790A and 1 μM A23187 in NRVMs treated with DMSO, PE, and PE plus GSK3874 (**G**). Quantification of [Ca^2+^]i response induced by GSK790A (**H**) and A23187(**I**)-induced in NRVMs treated with DMSO, PE, and PE plus GSK3874 (n = 10 per group). The arrow indicates the time of addition of GSK1016790A and A21387. All results represent mean ± SD, a one-way ANOVA followed by the Bonferroni test.

### TRPV4 Activation contributes to CaMKII phosphorylated

Ca^2+^/calmodulin-dependent protein kinase II (CaMKII) is upregulated after pressure overload and plays an essential role in cardiac hypertrophy and the progression of heart failure(Ljubojevic-Holzer, et al., 2020; Zhang, et al., 2003). More importantly, Ca^2+^ entry via TRPV4 can activate CaMKII in many other cells(Lyons, et al., 2017; Woolums, et al., 2020; Zhou, et al., 2021). Therefore, we hypothesized that TRPV4 activation contributes to cardiac hypertrophy through CaMKII. We first investigated the role of TRPV4 on CaMKII activation. Using NRVMs in vitro, we found that treatment with TRPV4 agonist GSK790A for 30 min markedly increases the expression of p-CaMKII (Thr287) compared with the DMSO group. However, GSK790A-induced activation of CaMKII was significantly blocked by either pretreating with TRPV4 antagonist GSK3874 or removing extracellular Ca^2+^, demonstrating that TRPV4-mediated Ca^2+^ influx promotes the activation of CaMKII (Figures 7A-D).

**Figure 7.**
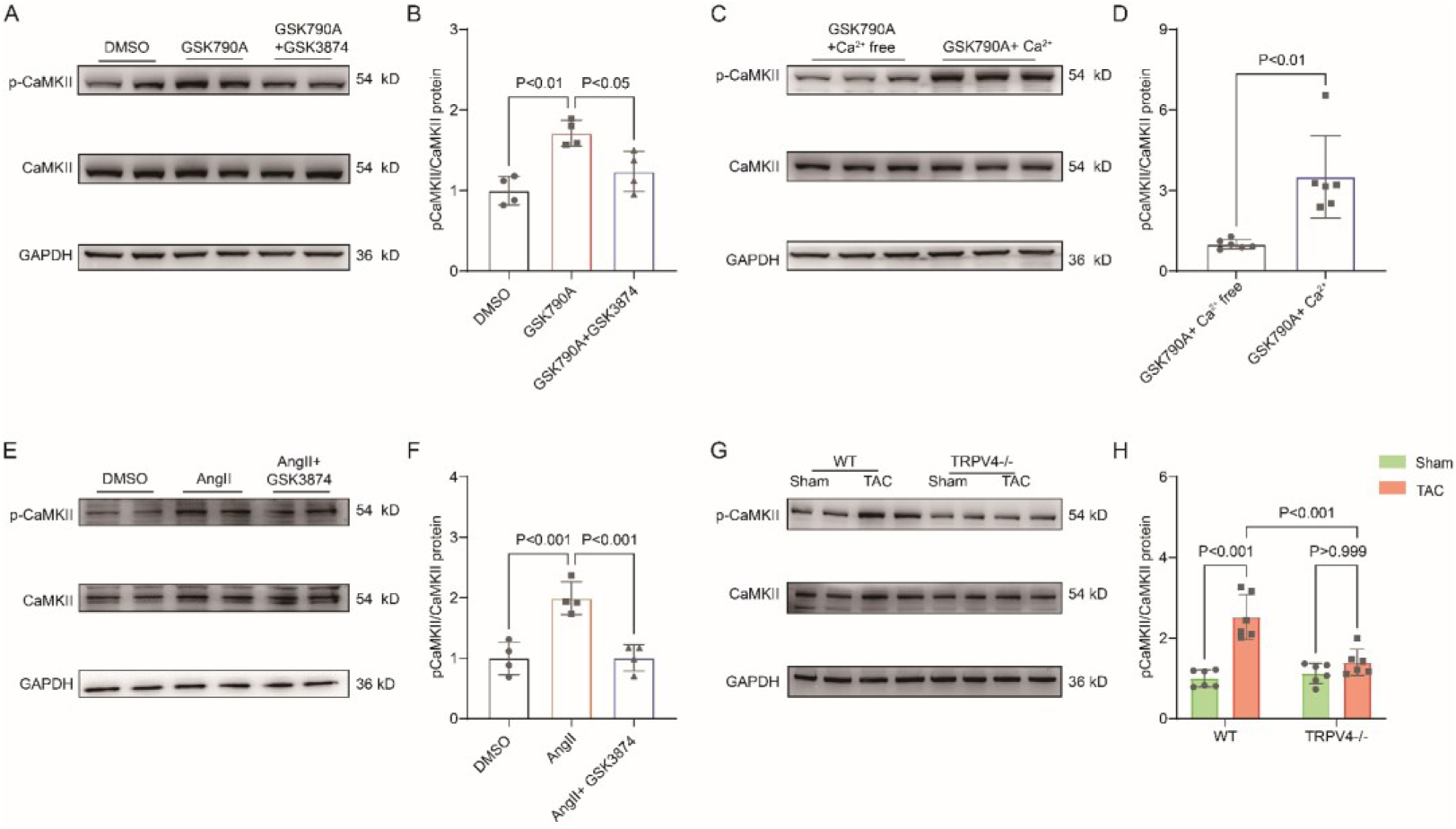
TRPV4 activation induces CaMKII phosphorylation. Representative immunoblot image (**A**) and statistics (**B**) of p-CaMKII/CaMKII in NRVMs treated with DMSO, GSK790A, and GSK790A plus GSK3874 (n = 4 per group). All results represent mean ± SD, a one-way ANOVA followed by the Bonferroni test. Representative immunoblot image (**C**) and statistics (**D**) of p-CaMKII/CaMKII in NRVMs treated with GSK790A in the absence and presence of Ca^2+^ medium (n = 6 per group). All results represent mean ± SD, an unpaired two-tailed student’s t-test. Representative immunoblot image (**E**) and statistics (**F**) of p-CaMKII/CaMKII in NRVMs treated with DMSO, AngII, and AngII plus GSK3874 (n = 4 per group). All results represent mean ± SD, a one-way ANOVA followed by the Bonferroni test. Representative immunoblot image (**G**) and statistics (**H**) of p-CaMKII/CaMKII in WT and TRPV4-/- mice at 4 weeks after sham or TAC operation (n = 6 per group). All results represent mean ± SD, a two-way ANOVA followed by the Bonferroni test.

Consistent with previous studies(Xiao, et al., 2018), NRVMs stimulated AngII for 48 h showed a 2-folds increase in p-CaMKII, and this response was largely abrogated by pretreatment with GSK3874 (Figures 7E-F). In vivo, TAC induced the upregulation of p-CaMKII in WT mice, but this response was not observed in TRPV4-/- mice (Figures 7G-H). Our results indicate that TRPV4 activation was required for the phosphorylation of CaMKII in response to pressure overload.

### TRPV4 activation promotes NFκB phosphorylation via a CaMKII-dependent manner

As shown in Figure 8A-B, a short-term (30 min) treatment with TRPV4 agonist GSK790A also dramatically increased the level of phosphorylated NFκB p65 in NRVMs. This effect was abolished by the pretreatment with TRPV4 antagonist GSK3874. Furthermore, AngII-induced phosphorylation of NFκB p65 was also prevented by pretreatment with GSK3874 (Figure 8C-D). Therefore, TRPV4 activation may promote the phosphorylation of NFκB p65.

**Figure 8.**
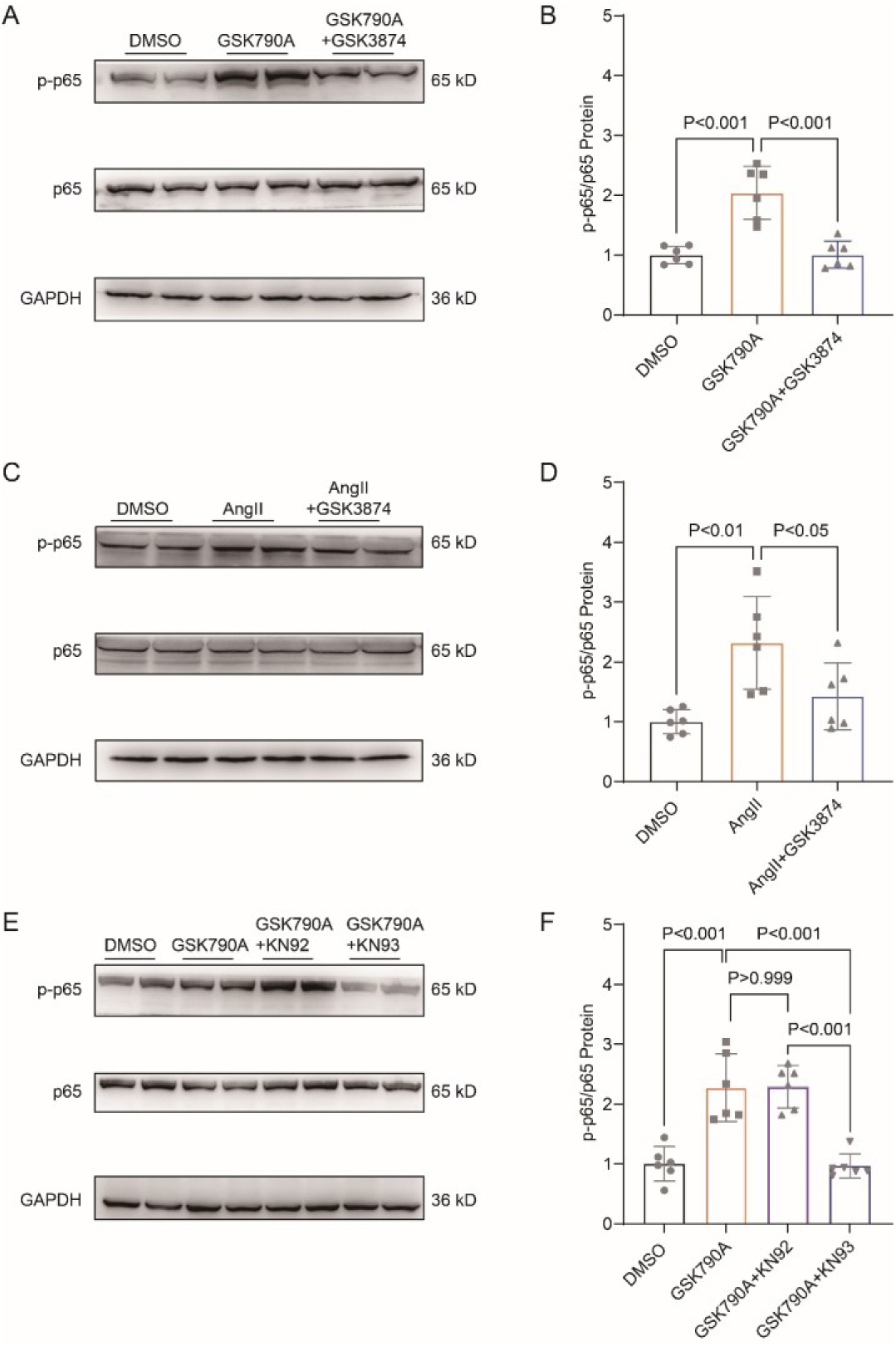
TRPV4 activation induces NFκB phosphorylation via a CaMKII signaling pathway. Representative immunoblot image (**A**) and statistics (**B**) of p-p65 /p65 in NRVMs treated with DMSO, GSK790A, and GSK790A plus GSK3874 (n = 6 per group). All results represent mean ± SD, a one-way ANOVA followed by the Bonferroni test. Representative immunoblot image (**C**) and statistics (**D**) of p-p65 /p65 in NRVMs treated DMSO, AngII, and AngII plus GSK3874 (n = 6 per group). All results represent mean ± SD, a one-way ANOVA followed by the Bonferroni test. Representative immunoblot image (**E**) and statistics (**F**) of p-p65 /p65 in NRVMs treated with DMSO, GSK790A, GSK790A plus KN92, and GSK790A plus KN93 (n = 6 per group). All results represent mean ± SD, a one-way ANOVA followed by the Bonferroni test.

We then asked how TRPV4 activation is linked to the NFκB signaling. Since the phosphorylation of NFκB could be regulated by the CaMKII signaling pathway(Ling, et al., 2013), we examined the involvement of CaMKII. Indeed, the application of a CaMKII inhibitor, KN93 (2 μM), abolished the GSK790A -stimulated NFκB p65 phosphorylation in NRVMs, supporting the role of CaMKII in linking TRPV4-mediated Ca^2+^ influx to NFκB activation.

## Discussion

In this study, we characterized the functional role of TRPV4 in pressure-induced cardiac hypertrophy and heart failure. We showed that TRPV4 activation promoted the development of pathological cardiac hypertrophy and heart failure. This effect was associated with Ca^2+^-mediated CaMKII phosphorylation and subsequently the activation of NFκB-NLRP3 (Figure 9). These results suggest that TRPV4 may be a potential therapeutic target for cardiac hypertrophy and heart failure.

**Figure 9.**
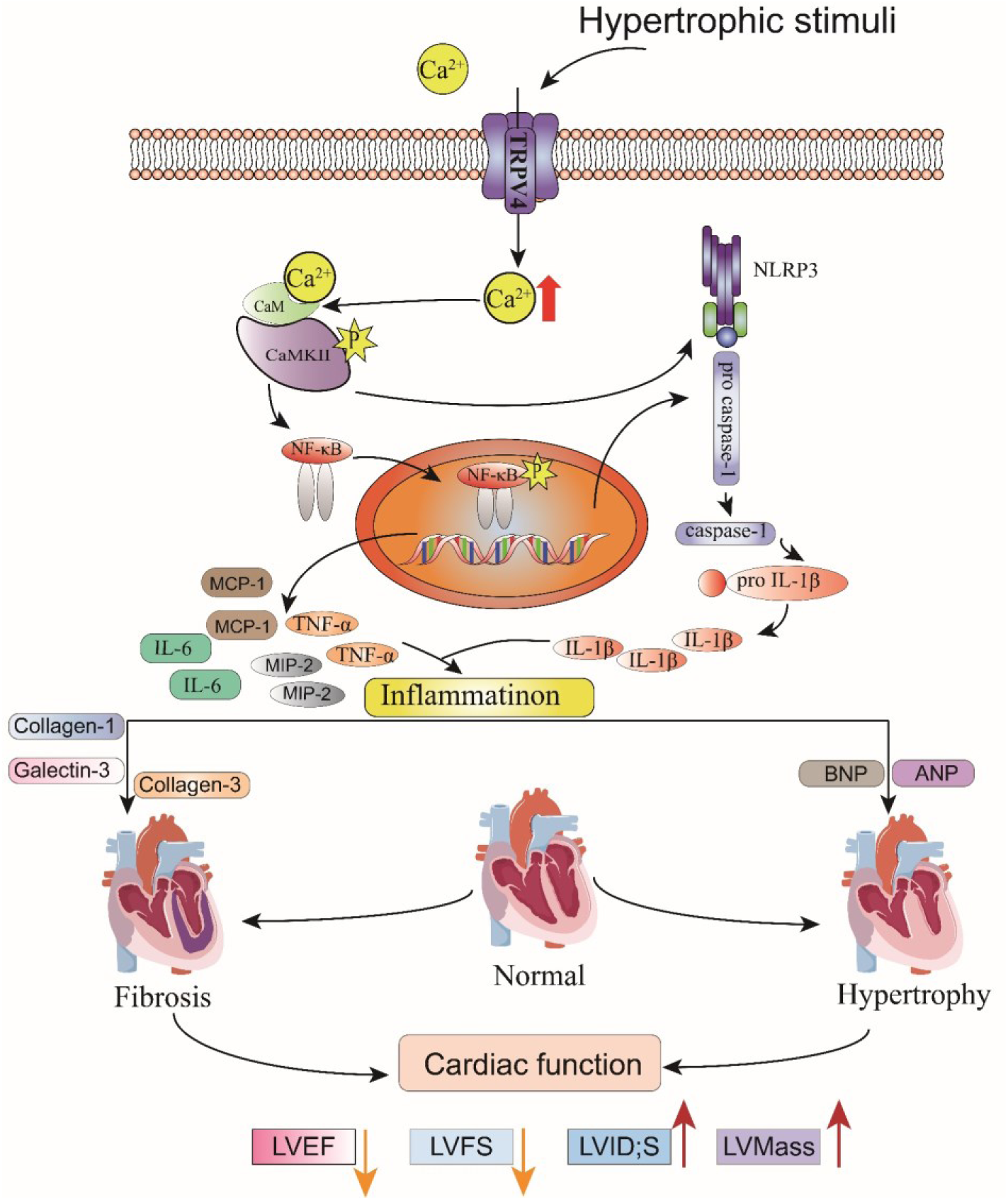
**Schematic illustration of potential mechanisms through which TRPV4 activation promotes pathological cardiac hypertrophy.**

As a non-selective calcium ion channel, TRPV4 is wildly expressed in the cardiovascular system and mediates cellular responses to a variety of environmental stimuli including hypo-osmolality, heat, and mechanical loading(Hof, et al., 2019; Randhawa and Jaggi, 2015). Previous studies including our own have demonstrated that TRPV4 is functionally expressed in hearts(Chaigne, et al., 2021; Peana, et al., 2021; Wu, et al., 2017) and can be upregulated by pressure overload(Morine, et al., 2016), following ischemia-reperfusion(Dong, et al., 2017; Jones, et al., 2019; Wu, et al., 2017), under inflammation conditions(Kumar, et al., 2020; Liao, et al., 2020), as well as after application of TRPV4 agonist GSK790A(Adapala, et al., 2016). Its activation induces Ca^2+^ influx and increases [Ca^2+^]i, which may subsequently promote cardiac remodeling and cardiac dysfunction. However, there are no data demonstrating the role of TRPV4 in pathological cardiac hypertrophy and heart failure in response to pressure overload. In the present study, we found that TRPV4 expression was significantly increased in mice hypertrophy hearts, human failing hearts, and AngII-induced hypertrophic cardiomyocytes, suggesting that TRPV4 was implicated in the processes of cardiac hypertrophy and failure. Furthermore, the deletion of TRPV4 attenuated TAC-induced cardiac hypertrophy and subsequence heart failure in vivo. Our in vitro experiments showed that TRPV4 blockade protected cardiac hypertrophy induced by AngII. Concomitant with this protection was the downregulation of multiple proteins and transcriptional markers associated with initiation and the progression of hypertrophy, inflammation, fibrosis, and heart failure. This data suggested that TRPV4 may play a role as either an initiating stressor or an upstream signaling transducer in response to pressure overload.

Recent studies have suggested that Ca^2+^ influx through TRPV4 can result in the activation of CaMKII(Lyons, et al., 2017; Woolums, et al., 2020). CaMKII can be rapidly activated in response to pressure overload and plays an essential role in cardiac hypertrophy and decompensation to heart failure(Baier, et al., 2020; Swaminathan, et al., 2012). Therefore, we hypothesized that TRPV4 experiences mechanical stress, mediates Ca^2+^ entry, and subsequently activates pro-hypertrophic signaling responses. Similar to our previous findings(Wu, et al., 2017), the TRPV4 agonist GSK790A induced robust Ca^2+^ entry in NRVMs. We also found GSK790A induced rapid phosphorylation of CaMKII, which could be prevented by TRPV4 antagonist and extracellular Ca^2+^ removal, demonstrating that Ca^2+^ entry following TRPV4 activation leads to CaMKII phosphorylation. Furthermore, AngII/PE-induced [Ca^2+^]i rise as well as the phosphorylation of CaMKII in NRVMs was significantly reduced by the TRPV4 antagonist. In addition, our in vivo studies showed that TAC-induced CaMKII phosphorylation was markedly blunted by genetic TRPV4 deletion. This evidence supports a key role of TRPV4 in mediating CaMKII activation during cardiac hypertrophy development.

Recent studies have shown that activation of CaMKII triggers NFκB-NLRP3 activation and leads to inflammation, which is important for the initiation and progression of pathological cardiac hypertrophy(Suetomi, et al., 2018; Willeford, et al., 2018). We found that TAC induced increases in IL-1β, IL-6, TNF-α, MIP-2, and MCP-1 expression, meanwhile the phosphorylation of p-65 and the expression of NLRP3, ASC, and cleaved caspase-1 were upregulated in WT mice. The above-enhanced effects, however, were diminished in TRPV4-/- mice. Similarly, AngII/PE-induced the upregulation of p-65 phosphorylation in NRVMs was reduced by the pretreatment with TRPV4 antagonist. These results suggest that TRPV4 activation promoted NFκB-NLRP3 activation and inflammation in response to pressure overload, which further demonstrated a mechanistic for TRPV4 in this response. Several other studies have also found that TRPV4 activation induces inflammation through the NFκB-NLRP3 signaling pathway(Wang, et al., 2021; Wang, et al., 2019). Additionally, we found that GSK790A also induced rapid phosphorylation of NFκB, which could be prevented by KN-93 for CaMKII inhibition, and this implies that CaMKII was involved in TRPV4 activation-induced the phosphorylation of NFκB. Therefore, our data continued to highlight the importance of TRPV4-mediated Ca^2+^ in intracellular signaling pathways and raise the possibility that TRPV4 activation promoted Ca^2+^ influx, led to the phosphorylation of CaMKII, and subsequently triggered the activation of NFκB-NLRP3, thus contributing to adverse cardiac remodeling.

An important limitation of our investigation is the use of the systemic functional abrogation TRPV4 model. TRPV4 is also expressed in cardiac fibroblasts and endothelial cells. Therefore, the effect of TRPV4 deletion on cardiac remodeling and dysfunction is not limited to cardiomyocytes. Interactions with cardiac fibroblasts or endothelial cells will need further study. Although the upregulation of TRPV4 was consistent in mouse hypertrophy hearts and human failing hearts, our data do not provide conclusive evidence about the involvement of TRPV4 in hypertensive cardiac damage in patients. Further human studies are needed to verify our results.

Collectively, our findings underscore the concept that TRPV4 might be a stress response molecule that is upregulated in cardiac hypertrophy. Activation of TRPV4 induced increases in Ca^2+^ influx, activated CaMKII, enhanced pro-inflammatory NFκB-NLRP3 signaling, promoted inflammation response, thus contributing to pathological cardiac remodeling. TRPV4 antagonism provides an exploitable therapeutic advantage for the treatment of cardiac hypertrophy and subsequent heart failure.

## Materials and methods

### Key resource table

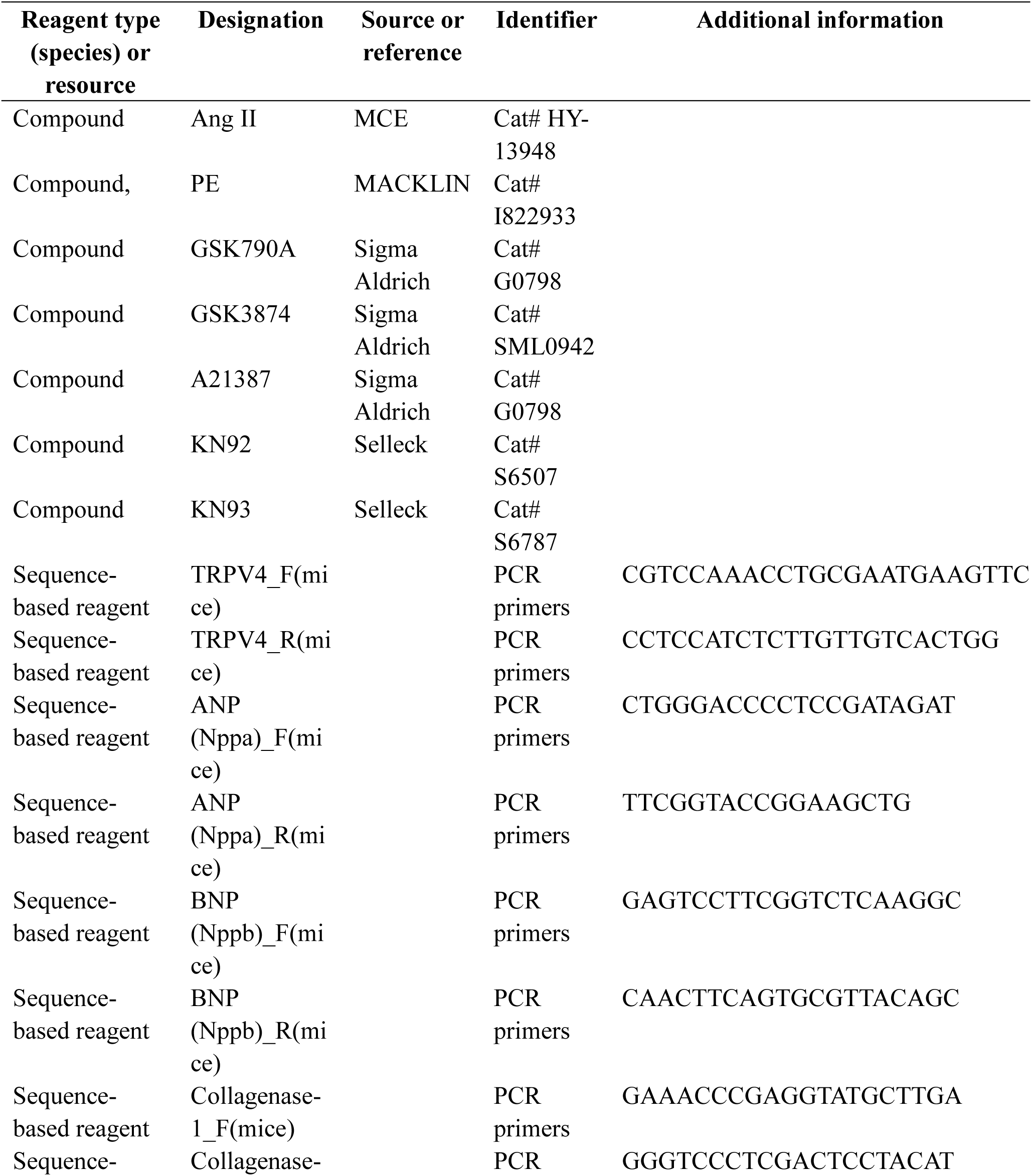
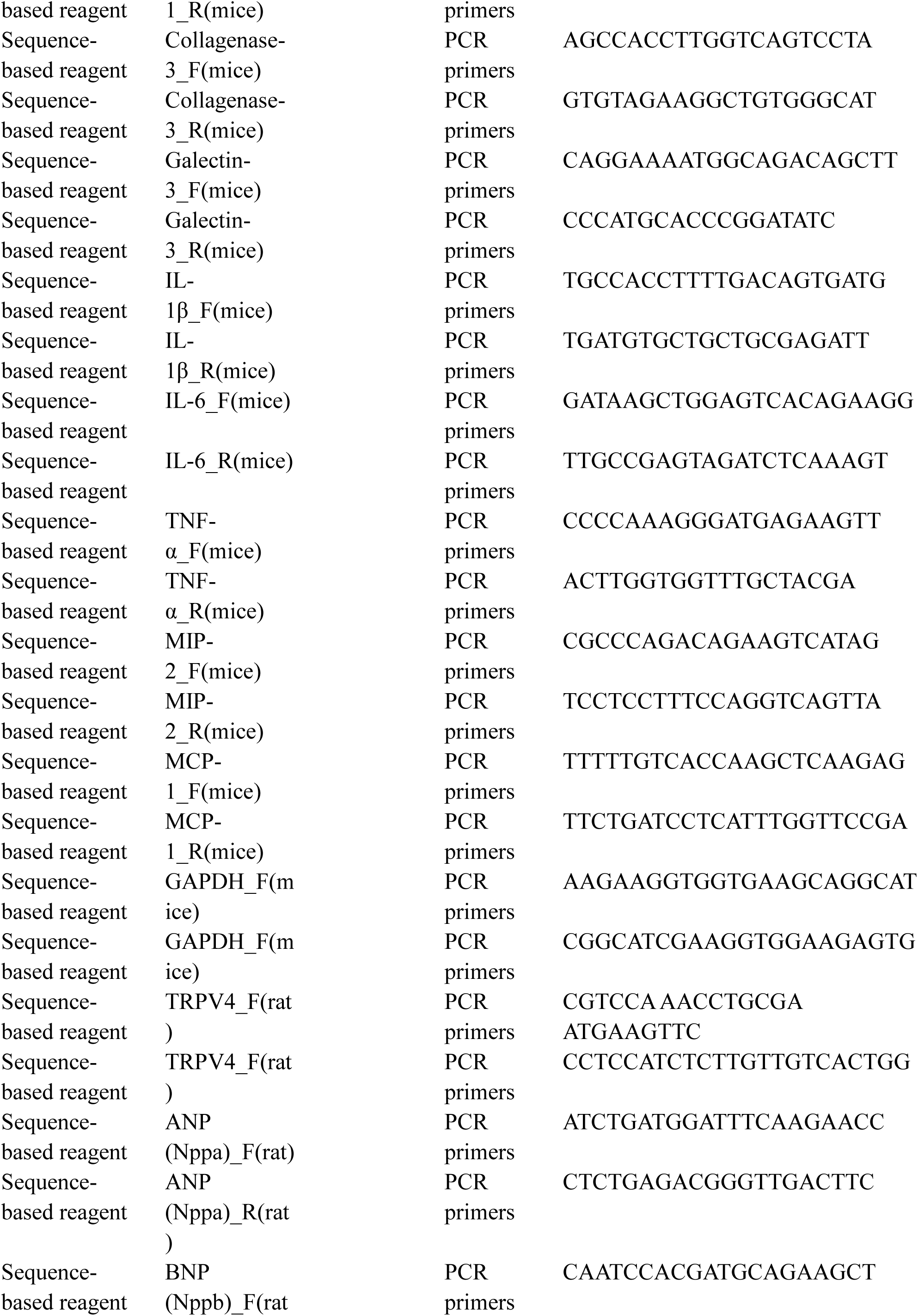
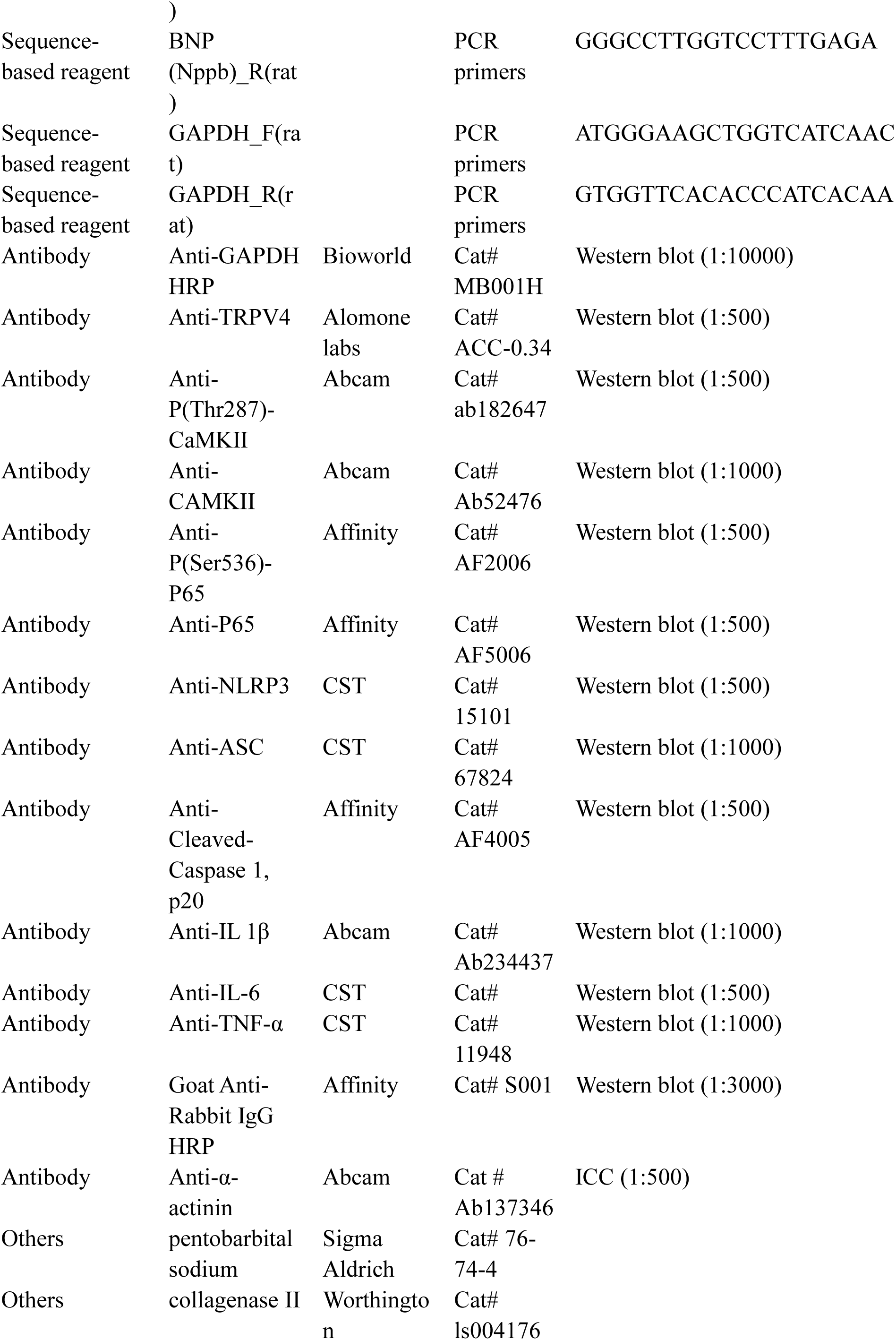
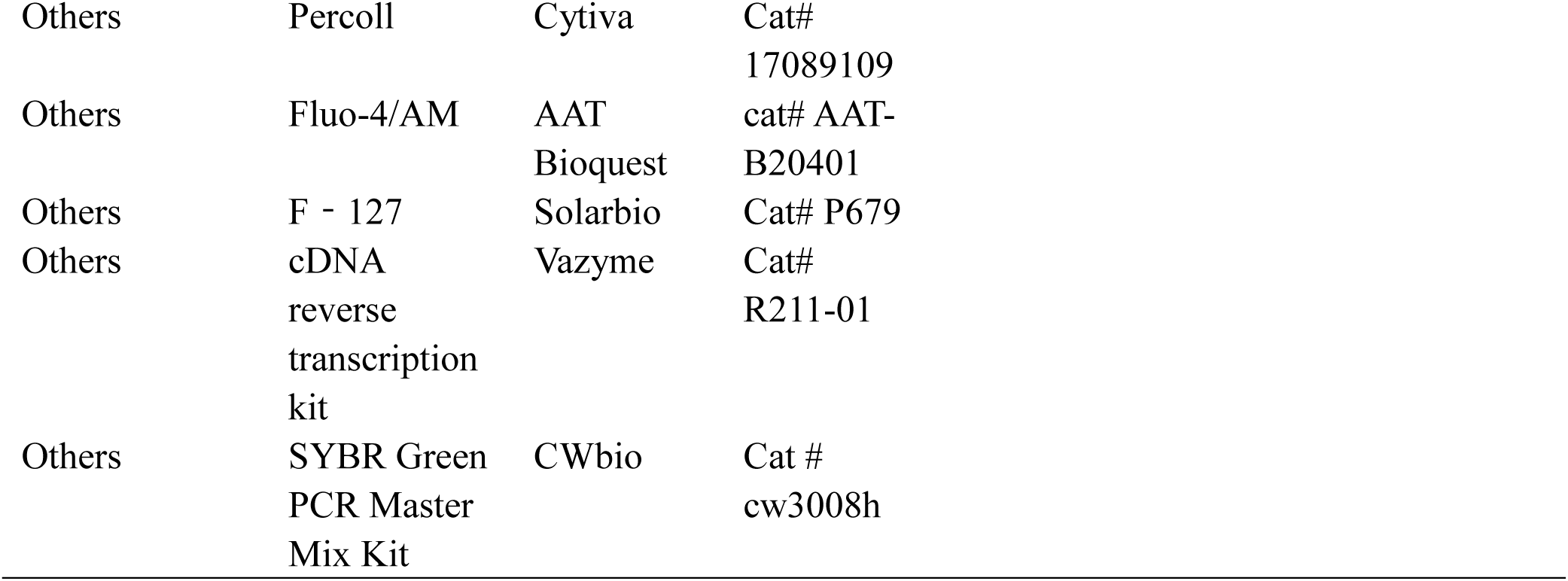

### Human heart tissues

Explanted, heart failure tissues were obtained from five patients with dilated cardiomyopathy (DCM) undergoing cardiac transplantation. Non-heart failure tissues were obtained from three organ donors whose hearts could not be placed due to size issues, ABO mismatch, or other factors. The study was in accordance with the Declaration of Helsinki (as revised in 2013). The study was reviewed and approved by the Ethics Committee of Union Hospital, Tongji Medical College, Huazhong University of Science and Technology (Wuhan, China; approval number: UHCT21001). Written informed consent was obtained from all the patients.

### Animals

Male C57BL/6 mice and new-born SD rats were purchased from the Laboratory Animal Center, Xuzhou Medical University (Xuzhou, China). TRPV4-/- mice were generated on C57BL/6 background as described previously(Dong, et al., 2017; Mizuno, et al., 2003). Genotyping was performed by PCR using ear punch/tail snip biopsies with the following primers: WT forward primer 5ʹ-TGTTCGGGTGGTTTGGCCAGGATAT-3ʹ and reverse primer 5ʹ-GGTGAACCAAAGGACACTTGCATAG-3ʹ, which produce a 796-bp product from the wild-type allele; knockout forward primer 5ʹ-GCTGCATACGCTTGATCCGGCTAC-3ʹ and reverse primer 5ʹ-TAAAGCACGAGG AAGCGGTCAGCC-3ʹ, which produce a 366-bp product from the target allele (Supplemental Figure S1A). RT-PCR of heart mRNA was used to confirm the deletion of TRPV4 sequence, indicated by a 534-bp cDNA fragment of WT mice, but absence in TRPV4-/- mice (Supplemental Figure S1B), as previously described (Boudaka, et al., 2020). All animal protocols were performed in adherence with the National Institutes of Health Guidelines and were approved by the Experimental Animal Ethics Committee of Xuzhou Medical University. Animals were housed in a temperature-regulated room (12 h day/12 h night cycle) with ad libitum access to food and water.

**Figure supplement 1.**
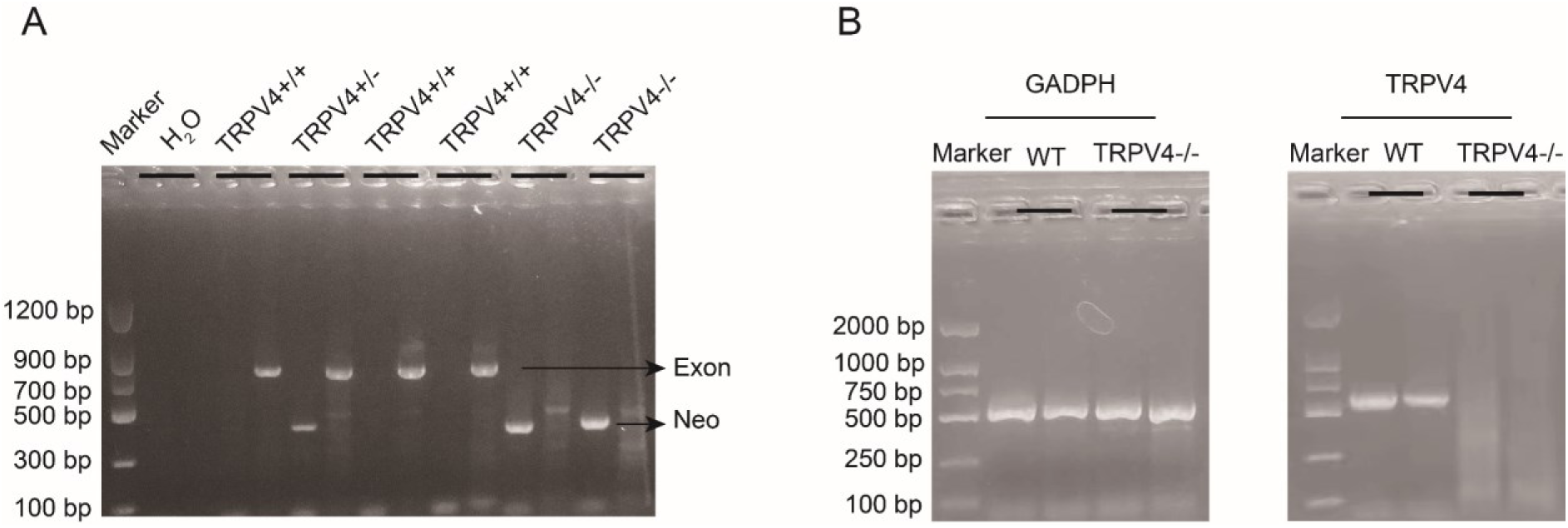
Genotyping of TRPV4 wild-type and TRPV4-/- mice and TRPV4 depletion in the heart of TRPV4-/- mice. A. Representative RT-PCR genotyping gel image of the WT, TRPV4+/-, and the TRPV4-/-. **B.** RT-PCR of total RNA from heart showing *TRPV4* mRNA was present in WT mice but absent in TRPV4-/- mice.

### TAC surgery

Eight- to 12-week-old male WT and TRPV4-/- mice were subjected to TAC to induce pressure overload. Mice were anesthetized by intraperitoneal (i.p.) injection of pentobarbital sodium (50 mg/kg), intubated via the oral cavity, and ventilated at 110 breaths/min. Following a sternotomy, the transverse aorta between the right innominate and left carotid arteries was dissected and banded with a blunt L type 27-gauge needle using a 5-0 silk suture. The needle was then removed. Successful TAC surgery was confirmed by measuring the right carotid/left carotid flow velocity ratio. The sham-operated mice underwent an identical procedure but without aortic constriction.

### Echocardiography

Echocardiography was performed 4 weeks after TAC by using a Vevo 2100 Ultrasound System (Visual Sonics, Toronto, Canada) as described in a previous study(Chen, et al., 2019). Briefly, the mice were anesthetized with isoflurane. Parasternal long- and short-axis views in B- and M-Mode were recorded when the heart rate of the mice was maintained at 430-480 beats/minute. The EF, FS, left ventricular end-systolic diameter (LVID), LV mass, and other function parameters were calculated with Vevo LAB software (Visual Sonics, Toronto, Canada) by a technician who was blinded to the treatment groups.

### Tissue collection

After the echo examination, the heart was harvested and rinsed with cold phosphate-buffered saline (PBS). After being weighted, the LV was cut into two parts. The top part was put into 4% paraformaldehyde for histological analysis, and the bottom part was quickly put into liquid nitrogen and transferred to a -80° freezer later. The HW normalized to BW and to TL were measured as indicators of cardiac hypertrophy(Zhao, et al., 2016).

### Histological analyses

For histological analysis, transverse LV sections were cut into 4-μm slices. The hematoxylin & eosin (H&E) staining was performed to analyze the histological change. Masson’s trichrome stain was performed to assess cardiac fibrosis. FITC-conjugated wheat germ agglutinin (WGA) was performed for further determination of cell size. A quantitative digital image analysis system (Image J software) was used in image measurement.

### Isolation of NRVMs and treatment

NRVMs were isolated according to previously established protocols(Golden, et al., 2012). In brief, LV from 1-3-day-old SD rats was harvested and digested in the presence of 0.5 mg/mL collagenase II at 37 °C. NRVMs were further purified by Percoll gradient centrifugation. Cells were plated at a density of 2.5 × 10^5^ cells/cm^2^ on collagen-coated plates and cultured in Dulbecco’s modified Eagle’s medium (DMEM) supplemented with 15% fetal bovine serum (FBS; Hyclone, USA), 100 units/ml penicillin, 100 μg/ml streptomycin, and 2 μg/ml cytosine arabinoside. The next morning, the media was changed to FBS-free DMEM for 24 h. Ventricular myocyte hypertrophy was induced by treatment with Ang II or PE for 48 h. In another group of experiments, cells were treated with TRPV4 agonist GSK790A (500 nM) according to the time required for the experiment, while TRPV4 antagonist GSK3874 (300 nM), KN92(2.0 μM), and KN93(2.0 μM) was applied 30 min earlier.

### Assessment of cell surface area

NRVMs were stained with antibodies for sarcomeric α-actinin and cell nuclei were counterstained with DAPI. Cell size was examined by TRICT-phalloidin staining assay and measured with Image J software.

### Calcium Fluorescence

Calcium imaging was performed as previously described(Wang, et al., 2019; Wu, et al., 2017). NRVMs were loaded with Fluo-4/AM (2 μM) and F-127(0.03%) for 30 min. Cells in 96-wells plates were illuminated at 488 nm and fluorescence emissions at 525 nm were captured by a multifunctional microplate reader (TECAN, Infinite® 200PRO, Swiss). Cells were stimulated with the TRPV4 agonist GSK790A (500 nM). A21387 (1 μM) was set as a positive control.

### RNA Extraction, cDNA Synthesis, and Quantitative PCR

Total RNA was extracted from LV tissues or cultured NRVMs using the Extraction Kit according to the manufacturer’s instructions. For cDNA synthesis, 500 ng RNA was reverse transcribed using a highly capacity cDNA reverse transcription kit. Real-time quantitative PCR (qPCR) was performed with SYBR Green PCR Master Mix Kit on a QuantStudio 3 system (Applied Biosystems, Foster City, CA). GAPDH was used as a housekeeper gene for the normalization of gene expression. The primers used in qPCR were listed in the Key resources table. The result for each gene was obtained from at least six independent experiments.

### Western blots

Total protein was extracted from LV tissues or cultured NRVMs with RIPA reagent. Then, protein expression was analyzed by standard western blot as described previously(Wu, et al., 2020). Briefly, protein (30 μg for each sample) was separated using 10% SDS-polyacrylamide gel electrophoresis and subsequently transferred onto polyvinylidene difluoride membranes (Millipore, Darmstadt, Germany). After 1 h of blocking with Western blocking buffer (CWbio, Taizhou, China), the membranes were incubated with primary antibody at 4 ℃. The next day, the membranes were washed with TBST and incubated with corresponding horseradish peroxidase (HRP)-conjugated secondary antibodies for 1 h at room temperature. Finally, proteins were visualized with the enhanced chemiluminescence kit (Affinity, Ancaster, ON, Canada). Band intensity was quantified by Tanon image plus software (Tanon, Nanjing, China). GAPDH was used as a loading control. The antibodies used in the study were listed in the Key resources table.

### Statistical analysis

All statistical data were presented as mean ± SD and analyzed by Graphpad prism 9. An unpaired two-tailed student’s t-test was used for comparison between the two groups. The differences among multiple groups were analyzed using one-way ANOVA or two-way ANOVA followed by the Bonferroni adjustment for multiple comparisons. *P*<0.05 was reported as statistically significant.

## Acknowledgements

The authors thank Prof. Atsuko Mizuno (Jichi Medical University, Japan) for TRPV4-/- mice. Dr. Yimei Du thanks Jenny Xiao (Columbia University, New York, USA) for editing the manuscript and checking for grammatical errors.

## Resource sharing

Further information and reasonable requests of resources and reagents should be directed to the lead contact, Yimei Du or Bing Han.

